# Topological Links in Predicted Protein Complex Structures Reveal Limitations of AlphaFold

**DOI:** 10.1101/2022.09.14.507881

**Authors:** Yingnan Hou, Tengyu Xie, Liuqing He, Liang Tao, Jing Huang

**Affiliations:** Key Laboratory of Structural Biology of Zhejiang Province, School of Life Sciences, Westlake University, 18 Shilongshan Road, Hangzhou 310024, Zhejiang, China; Westlake AI Therapeutics Lab, Westlake Laboratory of Life Sciences and Biomedicine, 18 Shilongshan Road, Hangzhou 310024, Zhejiang, China; Center for Infectious Disease Research, Westlake Laboratory of Life Sciences and Biomedicine, 18 Shilongshan Road, Hangzhou 310024, Zhejiang, China

**Author notes:** Corresponding author: Jing Huang.

**Keywords:** topological link, structure prediction, protein–protein interaction, AlphaFold, protein topology

## Abstract

AlphaFold is making great progress in protein structure prediction, not only for single-chain proteins but also for multi-chain protein complexes. When using AlphaFold-Multimer to predict protein–protein complexes, we observed some unusual structures in which chains are looped around each other to form topologically intertwining links at the interface. To our knowledge, such topological links are never observed in the experimental structures deposited in the Protein Data Bank (PDB). Although it is well known and has been well studied that protein structures may have topologically complex shapes such as knots and links, existing methods are hampered by the chain closure problem and show poor performance in identifying topologically linked structures in protein–protein complexes. Therefore, we address the chain closure problem by using sliding windows from a local perspective and propose an algorithm to measure the topological–geometric features that can be used to identify topologically linked structures. An application of the method to AlphaFold-Multimer-predicted protein complex structures finds that approximately 0.7% of the predicted structures contain topological links. The method presented in this work will facilitate the computational study of protein–protein interactions and help further improve the structural prediction of multi-chain protein complexes.

## Introduction

Predicting the three-dimensional (3D) structure of a protein from its primary sequence has long been a topic of great interest in biology. Recent studies using end-to-end deep neural network (DNN) methods such as AlphaFold [1, 2] and RoseTTAFold [3] have made great progress in this field. Reliable structure prediction of single-chain proteins has further inspired and facilitated the prediction of the structural details of protein–protein interactions (PPIs) [3, 4], which is pivotal for both the understanding of biological functions and intervention in diseases. Several recent studies have presented methods to predict the component identities and interaction modes of PPIs that are built upon AlphaFold [5–8].

Recently, we used AlphaFold-Multimer (v2.2.0) [4] to study PPIs and generated large-scale datasets of predicted protein–protein complex structures. We found that some of the predicted complex structures, including the top-ranked ones with the highest confidence scores, contained unusual topologically intertwining links at the interface (some examples are shown in Fig. 1). Intuitively, the formation of this kind of topologically intertwined interface in protein complex structures requires the unfolding of protein chains, which is nearly impossible under physiological conditions. Folded proteins encounter each other physically to form interacting complexes with conformational changes but do not unfold. In nature, topological links in protein complexes can be observed, but they always involve covalently modified amino acids or disulfide bonds, such as in the structures of virus capsids [9, 10]. We thus believe that the special topological links we observed in the predicted complex structures are likely artifacts generated from the prediction algorithm.

**Figure 1.**
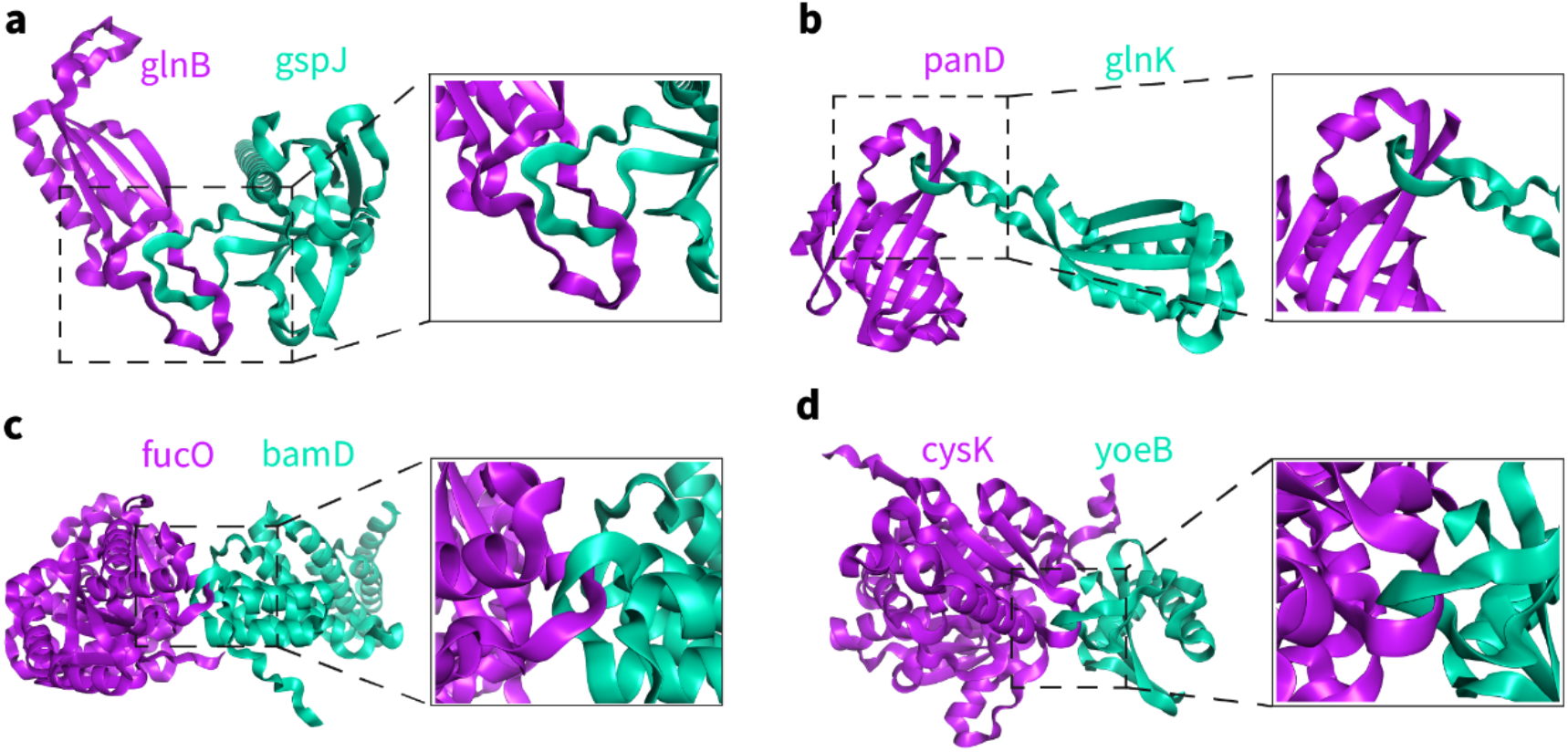
Topologically linked structures of protein–protein complexes in *E. coli* predicted by AlphaFold-Multimer (v2.2.0). The gene names of the proteins are marked. a) glnB-gspJ-4 (the fourth predicted structure ranked according to the confidence of AlphaFold-Multimer, same below), b) panD-glnK-10, c) fucO-bamD-1, d) cysK-yoeB-1.

Topologically complex elements such as knots, slipknots, and links have been well studied for single-chain protein structures [11–14], which are considered to be associated with particular thermodynamic and kinetic properties. Algorithms that identify and classify the topological features of these nontrivial protein structures are well established [15–22], and some researchers have attempted to apply them directly to multi-chain protein complexes. The identification of topological features for a set of closed curves is mathematically complete; however, protein chains are open curves with N- and C-termini such that determining how to close the curves represents a major difficulty. Existing strategies for closing a protein chain can be mainly classified into two categories: finding loops (e.g., finding covalent bonds and closing accordingly) [9, 10, 17, 23, 24] and creating loops (e.g., closing by connecting the N- and C-termini) [20]. The topological feature of a curve is deterministic, but it may vary according to different ways the curve is closed. Therefore, the key is to create properly closed loops in a protein chain without changing its original topological feature. To this end, approaches such as the KMT algorithm [13, 25], minimal surface analysis [26] and the Gauss linking integrals (GLN) method [27] are commonly used to determine the topological types and locations for protein structures with “closed” loops.

For topological link detection in multi-chain protein complex structures, determining how to close at least two curves without changing the original topological features is much more challenging. The LinkProt database [28] established a systematic classification of the topological links in protein structures with three categories: deterministic links, probabilistic links and macromolecular links. Deterministic links and macromolecular links are both topological links in structures that already have loops closed by covalent bonds (e.g., disulfide bonds); for such structures, there is no need to handle the chain closure problem. Probabilistic links are topological features in probabilistic form for structures with random closures, circumventing the chain closure problem. Another method of measuring the entanglement of protein complexes was established by observing the behavior of chains when pulling at both termini of each chain [29], where the closure of the chains and the results of identification are determined by the pulling directions. In addition, the Gauss discrete integral over open chains was suggested for measuring the entanglement in domain-swapped protein dimers without the need for closing curves [30], which is essentially an approximate estimation of the topological feature of two closed chains formed by directly connecting the N- and C-termini of each chain.

As the number of available experimental protein–protein complex structures is limited, existing methods have only focused on the global topological feature for the whole structure when detecting entanglement in multi-chain protein complexes. Currently, the explosively growing number of predicted structures of protein–protein complexes with real or fictitious interactions together with their more complicated interfaces bring new opportunities and challenges for the detection of topological features of protein complexes. We find that the existing methods may fail to identify many topological links observed in the AlphaFold-Multimer-predicted structures. Therefore, there is an urgent need for methods of identifying topological links in protein–protein complex structures in an accurate and robust manner.

To fill this gap, we propose a new method of detecting topologically linked structures in protein complexes by introducing geometric constraints when measuring topological features from a local perspective. Our method provides a solution to the chain closure problem based on the Gauss integral by systematically sliding windows in chains instead of focusing on the closures of the termini of chains, as in other studies. This is a fast and reliable method with deterministic results regarding how many topological links exist and where the links occur for a given protein–protein complex structure. We demonstrate how this method can detect topologically linked structures in several sets of predicted protein complex structures. Our work may further facilitate the improvement of protein structure predictions and computational PPI studies.

## Results

### Identification of topological links in protein complexes

To demonstrate why new algorithms are needed to identify topological links in protein–protein complex structures, we applied existing methods to 4 structures predicted by AlphaFold-Multimer v2.2.0 (Fig. 1) and 4 experimental complex structures from PDB (Fig. 2). For these four experimental structures, although two chains are entangled, it is evident that there are no topological links (Fig. 2). However, all 4 experimental structures are indiscriminately identified as topologically linked structures with LinkProt [28] and the GLN method presented in [17], as well as the pulling method [29]. These false inferences are caused by the closure of termini, which creates artificial topological links, indicating the low specificity of the link identification of these methods. For the predicted structures with topological links, LinkProt correctly identified three with Hopf links, but one (Fig. 1b, the 10th predicted structure of E. coli protein panD and glnK complex, named panD-glnK-10 hereafter) was unlinked (Table S1). A similar observation was made according to the close-to-zero whole GLN value (0.117). This false unlink inference was also caused by the closure of termini, which eliminated the original links; this happens particularly often for structures containing topological links formed with even-numbered windings in opposite directions in two chains, resulting in a cancellation effect when measuring the topological features from a global perspective. This is also demonstrated by the well-known failure in identifying the Whitehead link by existing methods [17].

**Figure 2.**
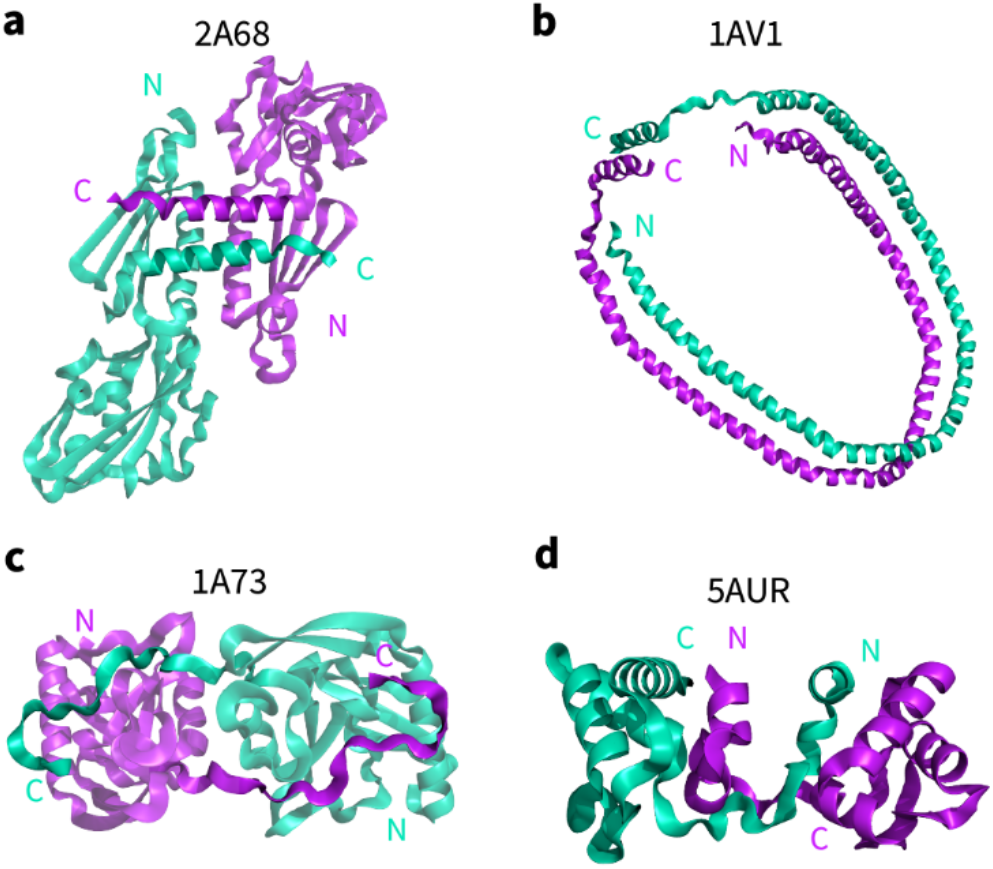
Experimental structures of protein–protein complexes in which the chains are wrapped but contain no topological links. The PDB codes are marked. N and C represent the N- and C-termini.

To overcome the chain closure problem and to detect topological links in protein–protein complex structures, we propose a novel algorithm by introducing an additional geometric dimension when systematically characterizing the topological features from a local perspective. As illustrated in Fig. 3, the algorithm is mainly composed of three steps: interface selection, systematic detection of atoms forming topological links and comprehensive inference. First, the protein chains are simplified by considering only the coordinates together with the peptide bonds of backbone N, Ca, and C atoms, which are represented by an ordered set of atoms. We further isolate the interaction interface by selecting atoms that are within a cutoff distance *D* =10 Å of each other chain, together with the atoms between them, forming two consecutive open subchains for analysis.

**Figure 3.**
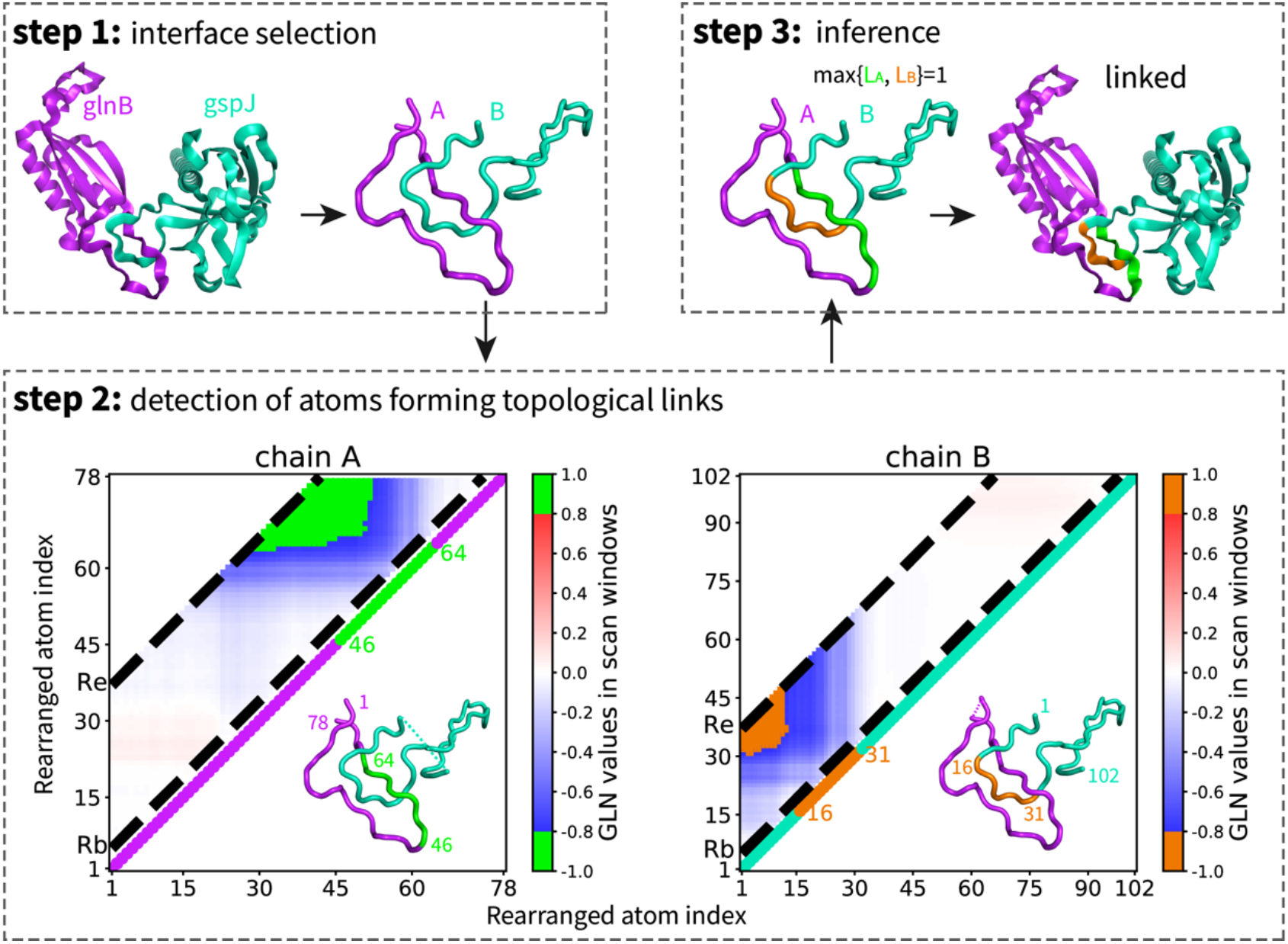
Schematic diagram of the algorithm for detecting topological links in protein–protein complex structures using the predicted glnB-gspJ-4 as an example (glnB as chain A in purple and gspJ as chain B in cyan). Step 1: Interface selection to preserve the topological relationships between chains. Each chain is replaced by its longest subchain whose backbone atoms at both ends are within *D*=10 Å of the other chain, and the atom indexes are rearranged. Step 2: The detection of atoms forming topological links by analyzing the GLN matrixes of both chains. For chain A (purple), each element in the GLN matrix (lower left panel) corresponds to the GLN value between chain B, closed at two ends, and a specifically closed fragment of chain A, where the index of the first atom is the row index and the index of the last atom is the column index. To systematically detect atoms forming topological links from a local perspective, we restrict the scanning windows to fragments whose lengths range between *R_b_*=4 and *R_e_*=36 atoms (the area between the two black dashed lines). Elements in the scanning windows with absolute GLN values over *T_s_*=0.8 are highlighted in green, indicating that the corresponding fragments form topological links with respect to chain B. The middle atoms of the highlighted corresponding fragments are marked to indicate the points at which topological links occur. The structure is shown in the lower triangular region, with the marked region (atom indexes from 46 to 64) also highlighted in green. Similarly, for chain B (cyan), the GLN matrix (lower right panel) is computed by closing chain A at its end points and sliding through different closed segments of chain B. The elements in the scanning windows with absolute GLN values over *T_s_*=0.8 are highlighted in orange, indicating where topological links occur. Step 3: The maximal number of fragments, with each formed by the marked adjacent atoms in chain A(*L_A_*) and chain B(*L_B_*), is used to measure the number of topological links (max{*L_A_, L_B_*}=1) in the structure, and the corresponding fragments in the two subchains are mapped back onto the original structure.

In the second step, we focus on the detection of atoms that form topological links in each chain.

We systematically create locally closed loops under geometric constraints by sliding windows along each chain to create fragments with varying lengths, in which each considered fragment is formed by several adjacent backbone atoms together with their peptide bonds and is closed by directly connecting its two ends. The topological feature of each fragment in the focused chain with respect to the other chain is measured by the GLN value, and the larger the absolute GLN value is, the more severe the entanglement of the two corresponding curves. All the fragments, each represented by its beginning and ending atoms, correspond to the form of a matrix; thus, the topological features can be measured by the GLN matrix constructed in this way. We restrict the systematic scan to fragments with lengths ranging from *R_b_*=4 to *R_e_*=36 atoms and evaluate the topological features according to the GLN values. Once the corresponding absolute GLN value of a fragment exceeds a threshold score (*T_s_*=0.8), indicating a possible topological link formed between the fragment and the other chain, the middle atom of the fragment, which contributes most to the topological link, will be selected and marked for further analysis in the next step.

In the third step, we infer the number of topological links of the protein complex structure with a comprehensive analysis of the marked atoms. Usually, the same topological link can be identified repeatedly around a set of marked atoms that have peptide bonds with each other, forming a consecutive fragment. Therefore, we calculate the number of consecutive fragments formed by the marked atoms in each chain and use the maximal one as the number of topological links in the structure, which is an indicator of whether a structure is topologically linked. In this way, the algorithm would not only identify whether the complex structure contains topological links but also locate the exact regions in which they occur. We note that the algorithm is robust with respect to the four user-definable hyperparameters (*D, T_s_, R_e_*, and *R_b_*).

Analysis of the 8 structures in Figs. 1 and 2 with the present method correctly found topological links for each of the 4 representative predicted structures and no topological links for any of the 4 deeply wrapped experimental structures (Table S1), demonstrating its superior ability to identify topological links in protein complex structures. We used 3 representative structures, glnB-gspJ-4 (Fig. 1a), panD-glnK-10 (Fig. 1b) and 2A68 (PDB code, chain A and B, Fig. 2b), as examples to illustrate why the addition of geometric constraints when characterizing topological features (termed topological-geometric features) from a local perspective makes a difference in identifying topological links. As shown in Fig. 4, glnB-gspJ-4 and 2A68 shared the same topological feature (Hopf link) from a global perspective, showing opposite topological-geometric features from a local perspective, and the cancellation effect of topological links from a global perspective in panD-glnK-10 could be avoided with a local perspective, indicating that topological–geometric features could provide additional key information to characterize protein structures. We also tested the method with the protein–protein docking benchmark set DB5.0 [31], and only one structure (PDB codes: 1NW9) was identified as containing topological links (Table S2). Manual inspection of this structure showed that one chain was broken at the interaction interface, so the presumably false positive was caused by too many missing residues at the interface (Fig. S1).

**Figure 4.**
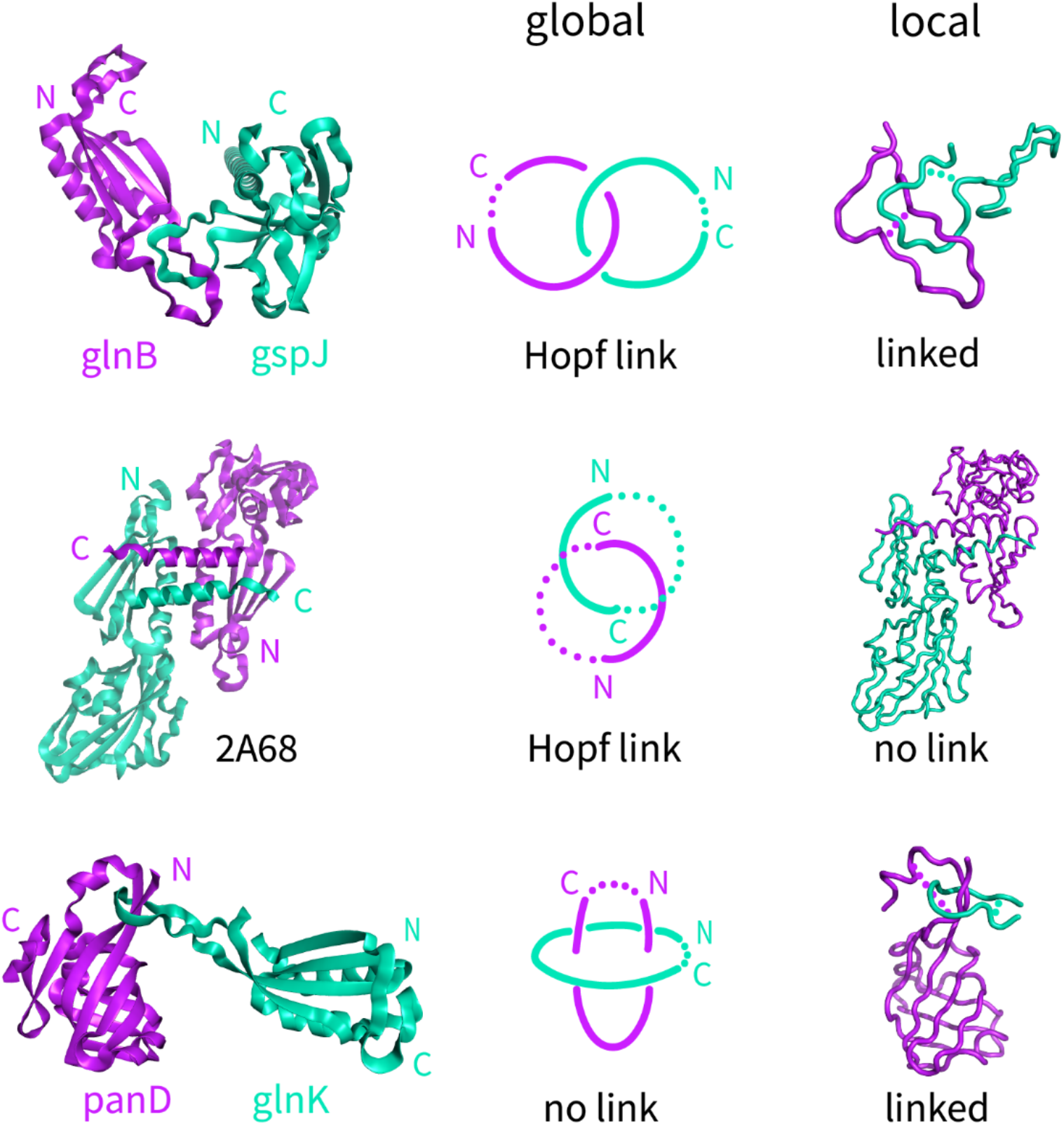
Comparison of the differences in characterizing topological features from global and local perspectives. The representative structures in the left column are two predicted structures (glnB-gspJ-4 and panD-glnK-10) and one experimental structure (PDB code 2A68, chain A in purple and chain B in cyan). The middle column represents the topological feature of protein complexes characterized from a global perspective in which two closed chains are formed by directly connecting the N- and C-termini of each chain. The right column represents the addition of geometric constraints to the topological features in the protein complexes from a local perspective, which create reasonable locally closed loops along each chain.

### Applications to protein complex structures predicted with AlphaFold-Multimer

Our algorithm allows quick identification of topologically entangled structures from a large number of predicted protein–protein complex structures. To quantify the number of topologically linked structures predicted by AlphaFold-Multimer (v2.2.0) from protein–protein complexes within the same genome, we generated a dataset by reorganizing protein–protein complexes from some known PPIs in the *Escherichia coli* protein interactome and predicting their structures with AlphaFold-Multimer. We randomly selected 20 PPIs from a benchmark set of experimental structures of the *E. coli* protein interactome in the PDB [32], split the two chains of each of the 20 PPIs into two categories (Table S3), constructed 400 (=20×20) paired protein complexes from the two categories and generated 10,000 predicted structures of the 400 protein complexes using AlphaFold-Multimer, with 25 predicted structures for each protein pair. We applied our method to this dataset and found that 60 structures contained topological links (Table 1 and Fig. S2). Focusing on the structures with the highest model confidence according to AlphaFold-Multimer, i.e., the top-ranked structures among the 25 predictions for each pair, 4 out of the 400 structures were identified as containing topologically linked structural elements (Fig. 1c-d, Fig. S2a-b).

**Table 1.** Summary of link detection by the proposed method on the four sets of AlphaFold-Multimer (v2.2.0) predicted structures.

| Species |  | All predicted structures |  |  | Top-ranked structures |  |  |
| --- | --- | --- | --- | --- | --- | --- | --- |
| Bait proteins | Prey proteins | Total | Linked | Percentage | Total | Linked | Percentage |
| *E. coli* | *E. coli* | 10000 | 60 | 0.60% | 400 | 4 | 1.00% |
| *Homo sapiens* | *Homo sapiens* | 18000 | 65 | 0.36% | 720 | 3 | 0.42% |
| *Homo sapiens* | SARS-CoV-2 | 14350 | 169 | 1.18% | 574 | 5 | 0.87% |
| Measles virus & Human<br>herpesvirus | *Homo sapiens* | 6425 | 58 | 0.90% | 257 | 3 | 1.17% |
| total |  | 48775 | 352 | 0.72% | 1951 | 15 | 0.77% |

We further tested our algorithm on three other sets of predicted protein–protein complex structures, among which one set concerns antibody–antigen interactions. Using 40 antibodies with known structures and 18 human interleukins (including IL2 and IL6, details in Table S3), we generated 18000 (40 × 18 × 25) predicted structures of antibody-interleukin complexes with AlphaFold-Multimer. To mimic practical applications in studying host–pathogen interactions, we used the nonstructural proteins (NSPs) of SARS-CoV-2 together with measles virus hemagglutinin glycoprotein (H) and human herpesvirus interleukin-6 homolog protein (vIL-6) to construct protein complexes with human proteins (details in Table S3), resulting in two datasets with 14350 and 6425 structures predicted by AlphaFold-Multimer. We note that in these datasets, most protein pairs will probably not interact under physiological conditions. Applying our method, topologically linked structures were identified in 65 out of 18000, 169 out of 14350, and 58 out of 6425 structures, respectively. For the top-ranked predictions with AlphaFold-Multimer, approximately 0.7% of structures were identified as topologically linked structures in all three sets (Table 1).

For these identified topologically linked structures of predicted antibody–interleukin complexes, we found that all the topological links occur between the antibody chains and the interleukin chain, while none of them occur between the heavy chain and the light chain of the antibody. This is probably due to the significantly larger number of heavy chain–light chain interactions in the training data of AlphaFold. We manually inspected the identified topologically linked structures, especially the top-ranked ones, and confirmed the existence of topological links based on the location information provided by the method (Fig. S3). In particular, we observed that more than half of the topological links occur around the complementarity-determining region (CDR) loops of the antibodies, as highlighted in orange in Fig. 5a-b and Fig. S3b for three representative structures. In addition, there are 2 predicted structures in which the interleukin chains form topological links with both the heavy chain and the light chain of the corresponding antibodies (Fig. S4). A recent study also pointed out the limitations of AlphaFold-Multimer in predicting antibody–antigen co-structures [33]. Inspection of the identified topologically linked structures in the other two datasets of virus and human protein complexes show similar results, and some exemplary structures are illustrated in Fig. 5c-d and Fig. S5-S6.

**Figure 5.**
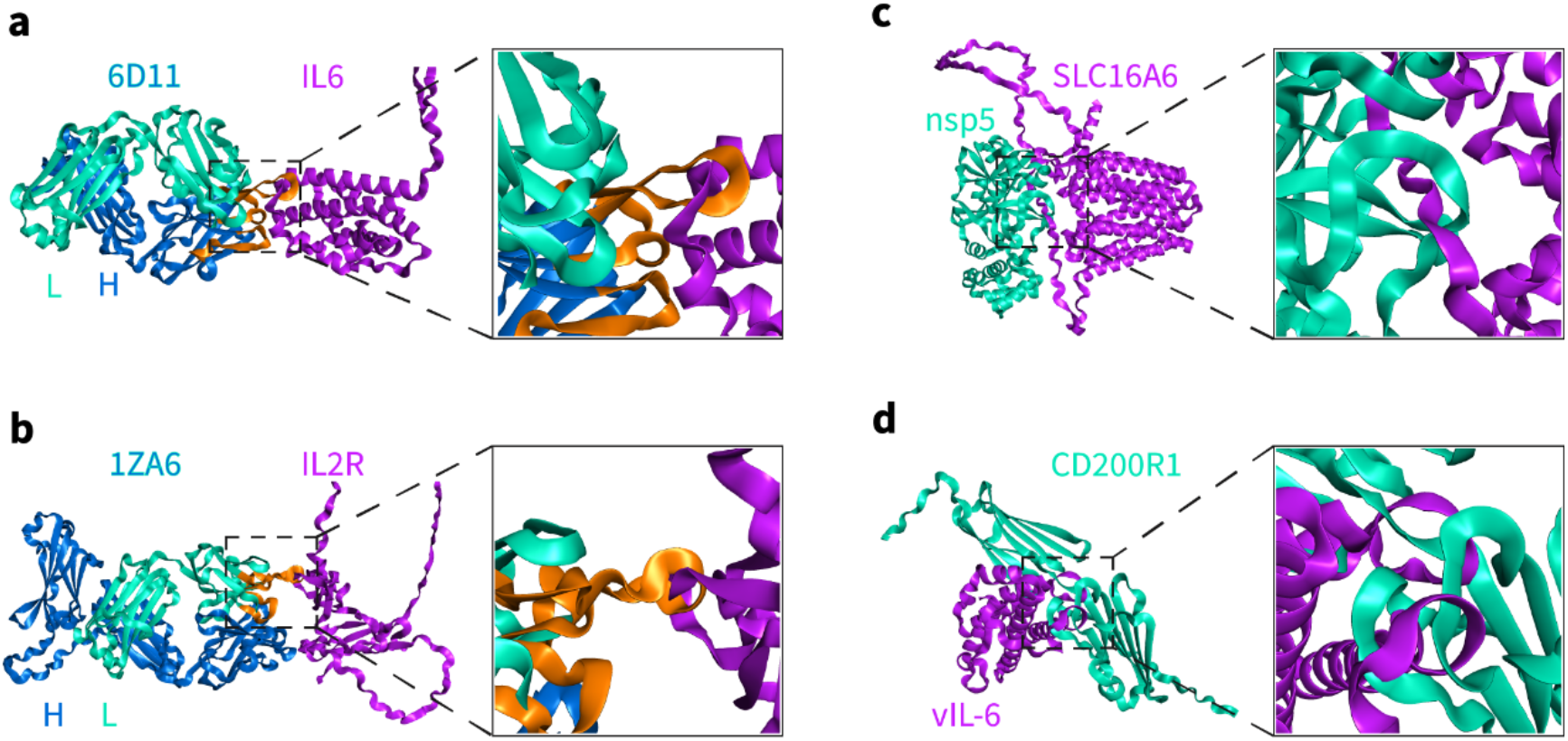
Topologically linked structures of protein–protein complexes predicted by AlphaFold-Multimer (v2.2.0). a-b) Two representative predicted structures (6D11-IL6-16 and 1ZA6-IL2R-1) of antibody-interleukin complexes in *Homo sapiens*. The CDR loops in the chain where topological links occur are colored orange. The PDB codes of the antibodies and the gene names of the interleukins are provided, with L indicating the light chain and H indicating the heavy chain. c) One representative predicted structure of complexes formed by human MFS transporters and NSPs of SARS-CoV-2 (SLC16A6-nsp5-1). d) One representative predicted complex structure of pathogenic virus and human membrane proteins (vIL-6-CD200R1-1).

## Discussion

Proteins physically interact with other proteins to function in many physiological processes, such as signal transduction and immune response. Recent advances in protein structure prediction have opened a new door to computationally determining the structures of protein–protein complexes using only the knowledge of their primary sequences. Currently, predicted protein structures are widely used by biologists. However, the booming number of predicted structures also raises new questions that need to be addressed. Clearly, evaluating predicted structures will be the next research focus after protein structure prediction. In this study, we focus on the topological links observed in predicted complex structures using the current version of AlphaFold-Multimer (v2.2.0) and then focus on detecting them. We developed an algorithm to identify topologically linked structures for protein complexes by providing a solution to the chain closure problem from a local perspective, that is, systematically sliding windows along each chain instead of closing the N- and C-termini. The algorithm has high sensitivity, as all the topologically linked structures it identified in this work were confirmed by manual inspection. The method runs in seconds for one complex structure, so it is computationally efficient and convenient for large-scale scanning of structures for topological links.

Topological links between subchains of the same chain can be formed during the folding process, while it may not be a physiological phenomenon that topological links form between different chains of protein–protein complexes. Therefore, we believe that the topologically linked structures of protein complexes predicted by AlphaFold-Multimer occur due to the algorithm’s failure to distinguish between inter-chain and intra-chain interactions, indicating an intrinsic flaw in the current AlphaFold algorithm that is unlikely to be eliminated through the selection of the most confident structure in the predictions. We do note that the confidence scores obtained by AlphaFold-Multimer are relatively low in regions where topological links occur (Fig. S7). This limitation might manifest itself in the prediction of super-large protein complexes and assemblies, and significantly compromise the prediction accuracy. Overcoming this limitation would also be necessary for the further development of computational methods to understand the dynamics and kinetics of protein–protein interactions.

In addition to the characterization of the topological features of complex structures, our method provides a measurement for evaluating complex structure prediction programs. For example, AlphaFold-Multimer v2.2.0 generated 0.7% topologically linked structures on average during the prediction of protein–protein complexes, while a previous version (v2.1.0) produced 20.4% topologically linked structures (Table S4), indicating a significant improvement in structure prediction for multi-chain protein complexes with an improvement in avoiding the clashes introduced in v2.2.0 [4]. Avoiding topological intertwining in the PPI interface is not trivial for structure prediction [34, 35]. The algorithm for capturing the topological–geometric features can be used as an additional loss function to constrain the feasible space in the deep learning network of AlphaFold-Multimer, which may be helpful in reducing the number of abnormal structures during prediction. We hope this research will further facilitate the improvement of protein structure predictions and computational PPI studies.

## Methods and Materials

### Algorithm for identifying topological links

#### Selection of the interactive interface

A protein complex with two chains (chain A and chain B) is taken as an example, as a complex composed of *M* (≥ 3) chains can be split into *M*(*M* – 1)/2 combinations of two-chain complexes. Only backbone heavy atoms (N, Ca and C) together with their peptide bonds are considered, and only atoms within a cutoff distance *D* of each other’s chain together with the atoms between them are preserved; their coordinates are used for further analysis with the indexes rearranged. The first 15 residues from either the N- or C-termini of each chain are removed to reduce the possible impact of their flexibility.

#### Detection of atoms that form topological links

To take a local perspective on topological links and to reduce the computational complexity, one chain (A) is systematically scanned locally in a sliding-window manner, and the other chain (B) is closed by directly connecting its two endpoints. Any fragment formed by adjacent atoms in chain A with a reasonable scanned window length is closed by directly connecting its two endpoints. Together with the simply closed chain B, the detection of the topological feature of the two closed curves is straightforward. The contribution of topological features from the fragment rather than the end-connected part can be represented by the GLN value; the larger its absolute value is, the more severe the winding of the fragment to the closed chain B. Fidelity in link identification is ensured by filtering out inappropriate connections of fragment ends, which can be achieved by setting a threshold score for the GLN values.

Let *N_1_* and *N_2_* represent the numbers of atoms in chain A and chain B, respectively. Let *S*^1^(∈ *R*^*N*_1_×*N*_1_^) represent the GLN matrix that measures all the fragments of chain A that wind around a closed chain B, where the element 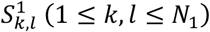 in the *k* th row and the *l*th column of *S*^1^ represents the GLN value of the fragment of chain A from the *l*th atom to the *k*th atom winding around chain B. Note that S^1^ is a lower triangular matrix, i.e., 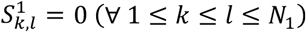. Similarly, *S*^2^(∈*R*^*N*_2_×*N*_2_^), representing the GLN matrix that measures all the fragments of chain B that wind around a closed chain A, will be analyzed in the same way.

To detect the atoms that form topological links, we focus on the fragments of the open chains whose lengths range from *R_b_* to *R_e_* (0 < *R_b_* < *R_e_* ≤ min{*N*_1_,*N*_2_}). Let 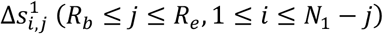 represent the absolute GLN values of the fragment from the *i*th atom to the (*i*+*j*) th atom in chain A, where 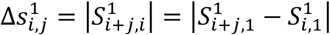. Here, we state that if 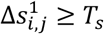, where *T_s_* is an empirical threshold score, then the ⌊*i* + *j*/2⌋ th atom, i.e., the middle atom of the corresponding fragment, is selected and marked because it may form topological links in chain A with respect to chain B. Note that one atom may be marked several times in fragments of different lengths.

#### Calculation of the topological link number

After a systematic scan of all the fragments in the length interval [*R_b_*, *R_e_*] in each chain, the same topological link can typically be identified repeatedly around a set of marked atoms, which are next to each other and form a consecutive fragment. Therefore, we calculate the number of consecutive fragments formed by the marked atoms in each chain and use the maximal one among all the chains to define the number of topological links in the structure of the protein complex. We call one structure a topologically linked structure if the number of topological links exceeds zero. In addition, the rearranged indexes of the marked atoms are mapped back to the whole structure, and the locations of the topological links in the form of residue indexes are returned by the algorithm.

There are four user-definable hyperparameters in the algorithm, i.e., the upper (*R_e_*) and lower (*R_b_*) limits of the length range, the threshold score (*T_s_*) and the cutoff distance between chains (*D*). The default values (*D* =10 Å, *T_s_* =0.8, *R_b_* =4 and *R_e_* =36) are empirically set without being elaborately crafted, which seems to be suitable for detecting topologically linked structures on the protein–protein interaction interfaces in this work. We do note that among these four hyperparameters, *R_e_*, which is the maximal fragment length in the scanning windows, is the most sensitive, so we smoothed the method by automatically adding 3 to *R_e_* for structures whose maximal absolute GLN value for the searched fragments was less than *T_s_* but greater than 0.9*T_s_*.

### Dataset generation

The protein sequences in this work were downloaded from the UniProt database, except that the sequences of the 40 antibodies were taken from the PDB database and the SARS-CoV-2 NSPs were taken from the NCBI database. Detailed sequence information is provided in the Supplementary Materials (Table S3). The predicted structures of the protein–protein complexes were generated by AlphaFold-Multimer (v2.2.0). For multiple sequence alignment (MSA) generation, we used UniRef90 v2019_10 (https://ftp.ebi.ac.uk/pub/databases/uniprot/previous_releases/release-2019_10/uniref/), BFD (https://bfd.mmseqs.com/), Uniclust30 v2018_08 (https://www.user.gwdg.de/~compbiol/uniclust/2018_08/), and MGnify clusters v.2018_12 (https://ftp.ebi.ac.uk/pub/databases/metagenomics/peptide_database/2018_12/). The sequence search tools were JackHMMER and HHblits (v3.3.0). For MSA pairing, we used UniProt (https://ftp.ebi.ac.uk/pub/databases/uniprot/current_release/knowledgebase/complete), downloaded on Feb 11, 2022. For the template search, we used “pdb_mmcif” from the PDB database as of Feb 10, 2022. The template search tools were hmmsearch and hmmbuild of HMMER (v3.3.2).

## Code availability

The source code has been deposited in a GitHub repository (https://github.com/JingHuangLab/topoLink).

## Acknowledgments

This work is supported by the National Natural Science Foundation of China (Grant No. 32171247, 21803057 to J.H., and 31970129 to L.T.), the Zhejiang Provincial Natural Science Foundation of China (Grant No. LR19B030001 to J.H. and LR20C010001 to L.T.), the Westlake Education Foundation, and Westlake Center for Genome Editing (Grant No. 20200000A992210/001 to L.T.). We thank the Westlake University Supercomputer Center for computational resources and related assistance.

