## Supplementary Material for "Topological Links in Predicted Protein Complex Structures Reveal Limitations of AlphaFold"

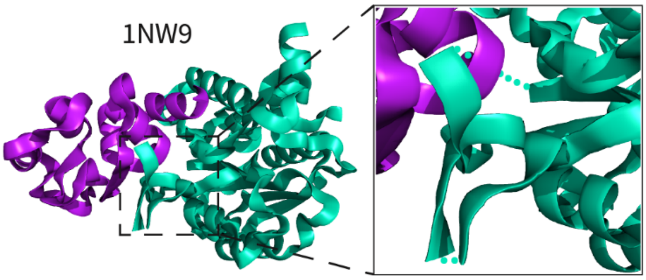


**Figure S1.** A broken chain (in cyan) at the interaction interface causes a topological link in the experimental structure (PDB code: 1NW9).

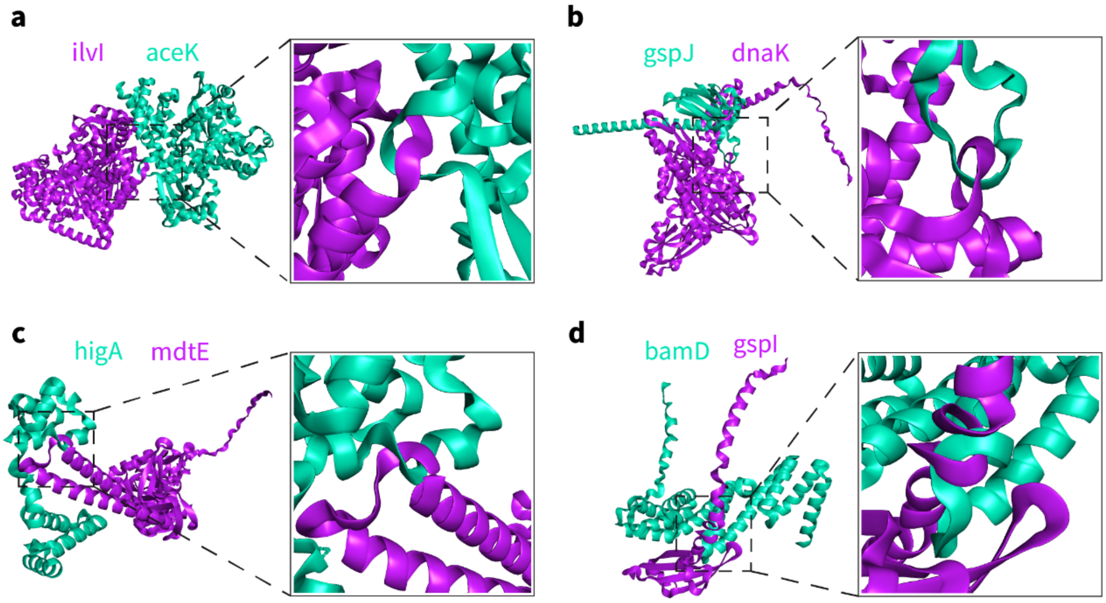


**Figure S2.** Topologically linked structures of protein–protein complexes in E. coli predicted by AlphaFold-Multimer (v2.2.0). The gene names of the proteins are marked. a) ilvl-aceK-1. b) dnaK-gspJ-1. c) mdtE-higA-2. d) gspI-bamD-2.

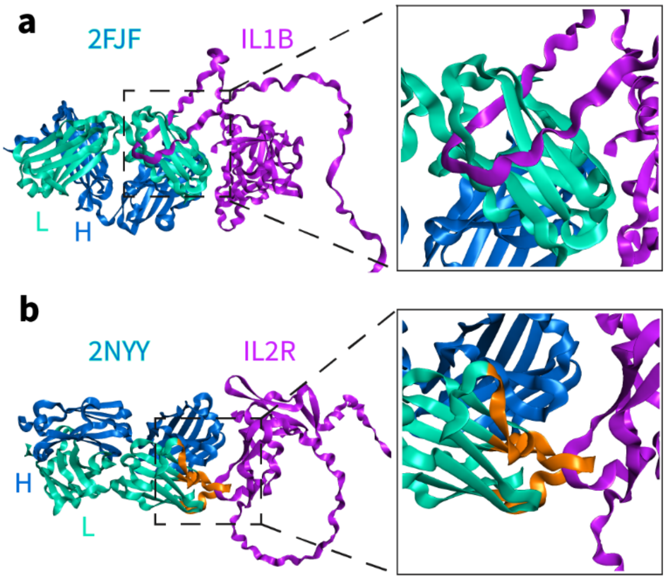


**Figure S3.** Topologically linked structures of antibody–interleukin complexes in *Homo sapiens* predicted by AlphaFold-Multimer (v2.2.0) with the highest confidence scores. The PDB codes of the antibodies and the gene names of the interleukins are provided, with L indicating a light chain and H indicating a heavy chain. a) 2FJF-IL1B-1. b) 2nyy-IL2R-1. The CDR loops in the chain where topological links occur are colored orange.

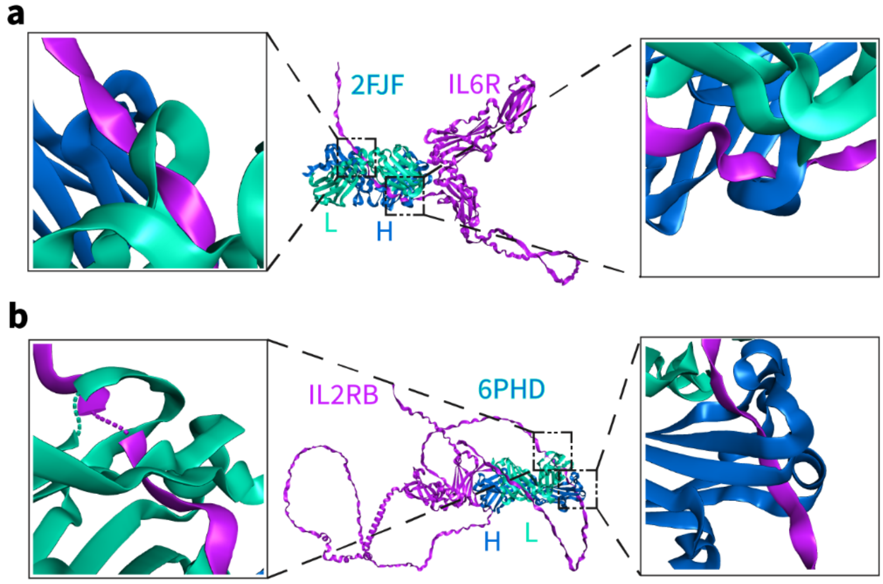


**Figure S4.** Topologically linked structures of antibody–interleukin complexes in *Homo sapiens* predicted by AlphaFold-Multimer (v2.2.0) with more than one topological link. The interleukins form topological links with both the heavy chain (H) and the light chain (L) of the antibodies. The PDB codes of the antibodies and the gene names of the interleukins are provided. a) 2FJF-IL6R-12. b) 6PHD-IL2RB-6.

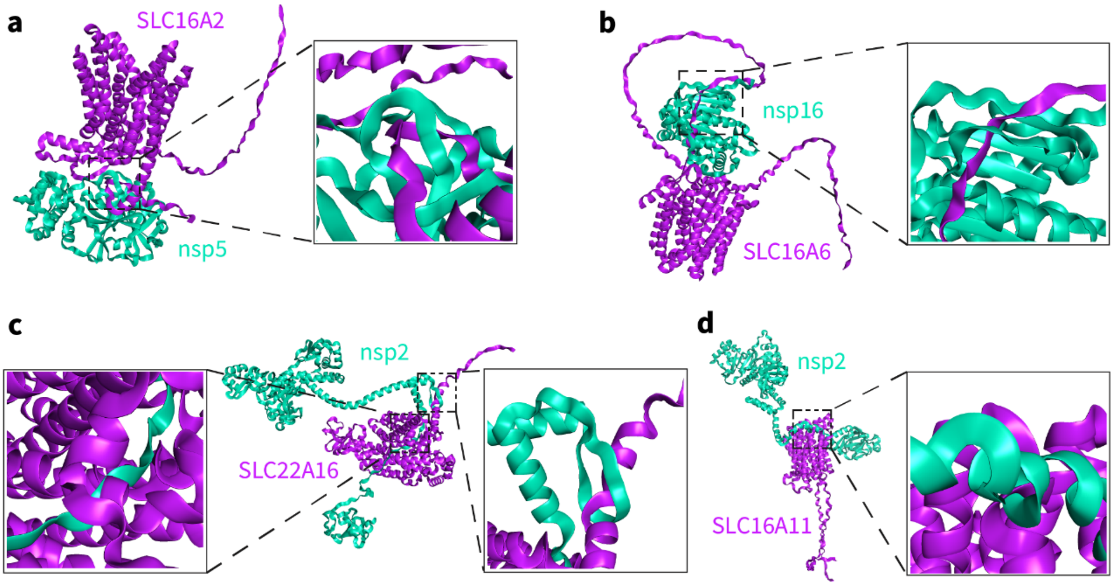


**Figure S5.** Topologically linked structures of complexes formed by human MFS transporters and NSPs of SARS-CoV-2 predicted by AlphaFold-Multimer (v2.2.0). The gene names of the human MFS transporters (purple) and the NSPs (cyan) are provided. a) SLC16A2-nsp5-1. b) SLC16A6-nsp16-1. c) SLC22A16-nsp2-1. d) SLC16A11-nsp2-1.

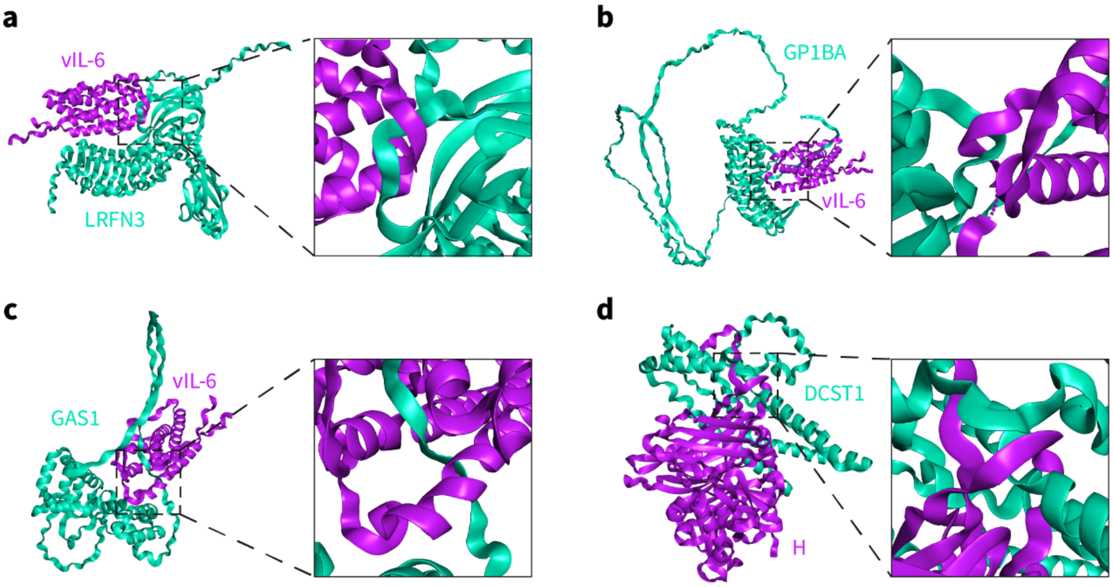


**Figure S6**. Topologically linked structures of pathogenic virus and human membrane protein complexes predicted by AlphaFold-Multimer (v2.2.0). The gene names of human herpesvirus interleukin-6 homolog protein (vIL-6, purple), measles virus hemagglutinin glycoprotein (H, purple) and human membrane proteins (cyan) are provided. a) vIL-6-LRFN3-1. b) vIL-6-GP1BA-1. c) vIL-6-GAS1-2. d) H-DCST1-14.

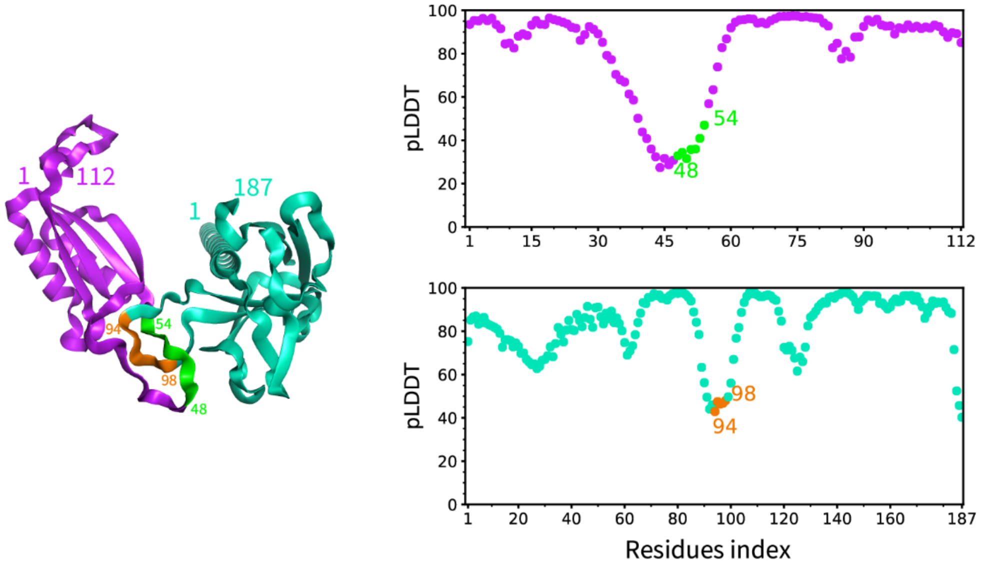


**Figure S7.** The confidence scores (pLDDT) of atoms provided by AlphaFold-Multimer (v2.2.0) are relatively low where topological links occur. An exemplary structure is glnB-gspJ-4.

**Table S1.** Comparison of methods for identifying topological links on the 4 AlphaFold-Multimer (AF2) [1] predicted structures and 4 experimental structures within labeled chains. Our method calculates and returns the numbers of topological links (NTLs) of these structures. The GLN method [2] was extended to apply to the 8 protein complexes, and the calculation was based on the GLN function in the topoly package [3]. Max|GLN| represents the maximal absolute GLN value between one chain and all fragments of the other chain, and whGLN represents the overall GLN value of the whole chain. The calculations with LinkProt were executed via its web server; for details regarding Hopf links with different subtypes depending on chirality and chain orientation, refer to [4].

| Structures | Source | Pair of chains | NTL | max\|GLN\| | whGLN | LinkProt | |
|---|---|---|---|---|---|---|---|
| | | | | | | type | probability |
| glnB-gspJ-4 | AF2 | A-B | 1 | 1 | -0.943 | Hopf.2 | 100% |
| panD-glnK-10 | AF2 | A-B | 1 | 0.973 | **0.117** | **unlink** | 100% |
| fucO-bamD-1 | AF2 | A-B | 1 | 1.149 | -1.031 | Hopf.2 | 100% |
| cysK-yoeB-1 | AF2 | A-B | 1 | 1.087 | 0.93 | Hopf.1 | 100% |
| 1A73 | PDB | A-B | 0 | 0.817 | 0.723 | Hopf.1 | 100% |
| 1AV1 | PDB | A-B | 0 | 0.839 | -0.713 | Hopf.2 | 100% |
| 2A68 | PDB | A-B | 0 | 0.934 | 0.934 | Hopf.1 | 100% |
| 5AUR | PDB | A-C | 0 | 0.813 | 0.754 | Hopf.1 | 100% |

**Table S2.** The detection of topological links in the experimental structures from the protein–protein docking benchmark DB5.0 set [5]. The table is available in Excel format as Dataset S2.

**Table S3.** Information on the proteins in the datasets generated in this work. The table is available in Excel format as Dataset S3.

**Table S4.** Summary of link detection by our method on the datasets of AlphaFold-Multimer (v2.1.0) predicted structures. Note that AlphaFold-Multimer generates 5 predictions for each protein pair.

| Species | | All predicted structures | | | Top-ranked structures | | |
|---|---|---|---|---|---|---|---|
| Bait protein | Prey protein | Total | Linked | Percentage | Total | Linked | Percentag e |
| *E. coli* | *E. coli* | 2000 | 227 | 11.35% | 400 | 26 | 6.50% |
| *Homo sapiens* | SARS-CoV-2 | 2400 | 1193 | 49.71% | 480 | 182 | 37.92% |
| Human herpesvirus | *Homo sapiens* | 1060 | 135 | 12.74% | 212 | 15 | 7.08% |
| total | | 5460 | 1555 | 28.48% | 1092 | 223 | 20.42% |
